# Nutritional status and morbidity profile of children with contact to leprosy in the rural community

**DOI:** 10.1101/452995

## Abstract

**Introduction:** Childhood leprosy reflects upon the disease transmission in the community. So, this study aimed to find out the children with contact to leprosy in their surroundings, prevalence of leprosy or subclinical infections in them and to assess their nutritional status. The study was conducted for 2 months and analysed 70 children in the rural community who were living with a household contact of leprosy.

**Methods:** 70 children in the rural areas surrounding Chengalpattu living with leprosy contacts were selected for carrying out the study. Information regarding their demographic characteristics, socioeconomic factors, environmental conditions, feeding practices, food habits and any present health problems or in the recent past were collected. The children were then subjected to anthropometric measurements. The children were clinically evaluated by a dermatologist qualified in paediatric leprosy and children who were diagnosed as cases of leprosy were classified according to Ridley-Jopling classification. Slit skin smears for acid fast bacilli was done in all children with suspicious skin lesions.

**Results:** Out of the 70 children taken into the study, 41 were boys and 29 were girls. 7-22% of boys and 3-6% of girls and overall,4-15% children are severely malnourished. 19 out of the 70 children had clinical pallor. Among the 70 leprosy contact children, 3 children were diagnosed to have leprosy (4.28%).Of the 3, 2 children had multibacillary leprosy while 1 had paucibacillary leprosy, according to the WHO classification and all 3 were classified as cases of Borederline Tuberculoid Leprosy according to Ridley-Jopling classification. All these 3 children had contact to leprosy for 10 or more years living with them.

**Conclusion:** It can be concluded that malnutrition, the closeness and duration of contact to leprosy are significant risk factors for leprosy. Regular contact screening and early case detection are essential strategies to prevent further transmission in the endemic areas. Diagnostic methods for detection of subclinical infection in contacts needs further research.

## Introduction

Leprosy, which is one of the earliest ever diseases known to mankind, is a public health problem in many developing countries. It is a disease which predominantly affects the skin and the peripheral nerves. Childhood leprosy reflects upon the disease transmission in the community and efficiency of various leprosy control programmes. Leprosy is still a significant health problem, even though its prevalence is decreasing in India. Childhood leprosy indicates endemicity of leprosy. **^1^**

The detection of new cases throughout the world isn’t declining over the years. India forms one of the major contributions to the worldwide leprosy case load. There are 16 high-burden countries from which 50% of the new leprosy cases in the world are reported, one of which is India. One of the risk factors for leprosy is the presence of an index case as a household contact. Early detection of cases is being emphasized by all health care professionals but active contact screening isn’t yet gaining that much importance. Active contact screening can prove to be an efficient tool to reduce the leprosy burden. **^2^**

Leprosy has been eliminated (prevalence less than 1 per 10000 population) from most of the countries, according to the World Health Organisation. India achieved elimination in December 2005. In India, there has been decrease of prevalence and new case detection rates but still children form about 10% of new cases detected. This shows there is still active transmission of M. leprae among Indian communities building endemicity. Children, in a household with a newly diagnosed leprosy case, are at increased risk, compared with the general population and even compared with adults in the household, to suffer from the disease. There is much higher risk to children aged 1–14 years than in older people. **^3^**

Leprosy in under 15 years old children reflects recent disease and active foci of transmission in the community.The prevalence of previously undiagnosed leprosy in the general population, in highly endemic areas, is six times higher than the registered prevalence. **^4^**

It is necessary to know that this disease can occur in children under 15 years of age, the risk being increased when they have contact to leprosy in their surroundings. Paediatricians and dermatologists should consider leprosy within all differential diagnoses, while providing medical care to children and adolescents. So the early diagnosis of leprosy is essential in the prevention of deformities, whose consequences are more catastrophic in children under 15 years of age. **^5^**

Leprosy has always been associated with the social stigma that comes along with it. It is mainly the disabilities which lead to leprosy patients being stigmatized and secluded from general public. Disabilties in children can have more serious effects than adults. Disabilities can affect children occupationally which can affect them economically in their future. **^6^**

Moreover the various risk factors associated with leprosy patients and their contacts are still not clearly known. Only through intensive research helps in examination of group-level and individual level factors. This study hence purported to detect possible risk factors for development of disease in leprosy contact. **^7^**

Even though there have been lots of research regarding childhood leprosy and children with contact to leprosy, there isn’t much research which concentrate on the nutritional status of children who are leprosy contacts. So, this study aimed to find out the children with contact to leprosy in their surroundings, prevalence of leprosy or subclinical infections in them and to assess their nutritional status.

It was started as an observational pilot study to analyse these parameters in such children. The study was conducted for 2 months and analysed 70 children in the rural community who were living with a household contact of leprosy.

## Aims and Objectives

The main issues to be examined are:

a. What is the prevalence, of children with leprosy contacts, in the rural community?
b. How is the nourishment profile of such children?
c. What could be the probable risk factors that can lead to infection in these children?
d. If transmission to children is still occurring, largely from undiagnosed cases in the community, can it be interrupted?
e. Can progression to leprosy disease be prevented in those contacts more exposed to infection or even in those already infected, but currently in a subclinical state?

## Methodology

### Study Design and Type of Study

This study was conducted as an observational pilot study involving the under 15 years old children, in the rural community, who have a household contact of leprosy, and hence there was no pre-determined sample size. A previously diagnosed case of leprosy living together with family members and sharing the same roof and meal from common kitchen is called a household contact. Children already diagnosed with leprosy were excluded. The study was conducted from August to September 2017.

### Study Methods

The cases recorded in Central Leprosy Teaching and Research Institute at Chengalpattu, in Kancheepuram district in the state of Tamil Nadu in India, who had children living with them in their homes, in the rural areas surrounding the institute, were selected. 70 such children were selected for carrying out the study. Selected children and their parents were briefed about leprosy and the main objectives of the study. Informed written consent was obtained from the parents or another responsible adult within the family of the children, accompanying them (since the study involved under 15 years old children), who agreed to take part in the study.

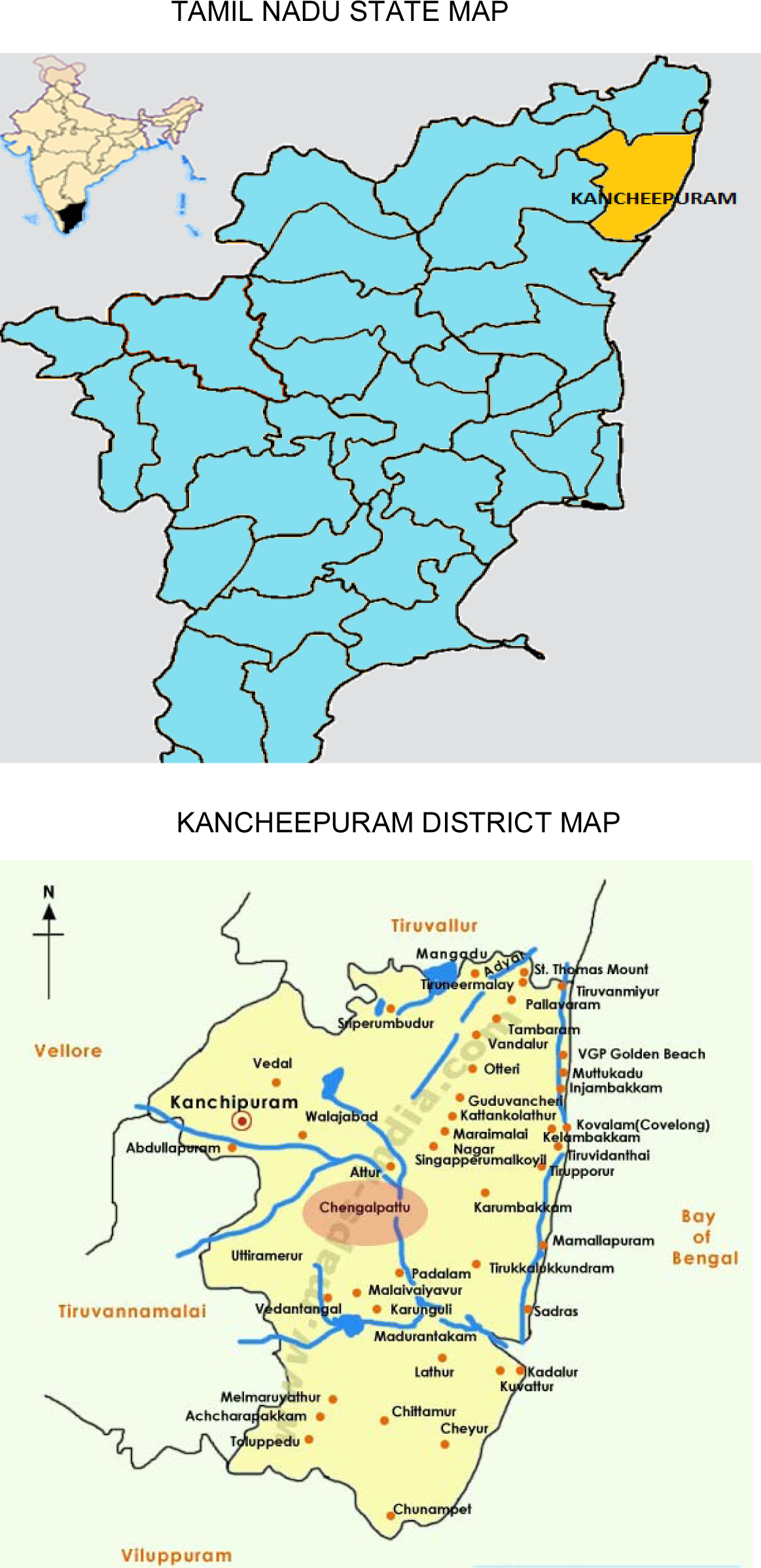

### Nutritional Status Assessment

One to one interview method was employed to collect information regarding demographic characteristics, socioeconomic factors, environmental conditions, feeding practices, food habits and any present health problems or in the recent past. The children were then subjected to anthropometric measurements;

- body weight measurement (to the nearest kg) with the subject standing on the weighing scale as per WHO guidelines.
- height measurement (to the nearest cm) with the subject standing in erect position against a stadiameter for all children.
- BMI was calculated using this formula: BMI = Weight (in kg)/ Height (in m^2^)
- By calculating BMI, the nutritional status of the children was assessed. The mean value of BMI at different age points was compared with the corresponding reference values of National Health and Statistics Report. **^8^**
- Head circumference measurement (for 0-5 years children) with a measuring tape placed over the occipito-frontal circumference, just over the eyebrows and occiput
- Mid arm circumference measurement (for 0-5 years children) of the left arm at the midpoint between acromion process and olecranon process, with arm lying laterally to the trunk. **^9^**

### Clinical evaluation

The children were clinically evaluated by a dermatologist qualified in paediatric leprosy. General examination and then specific examinations of skin, eyes, ear, nose, throat, tooth, CVS, Chest, Abdomen and CNS was done. In every child, a detailed clinical examination of the children was carried out for any skin lesions, with sensory loss in them. Girls were assessed by a female examiner. In case of any suspicious skin lesion, they were noted down and further testing was done.

Nerve function assessment was done by Voluntary Muscle Testing (VMT) for motor function of muscles supplied by the nerve. Sensory testing for sensory loss was done in the areas supplied by the nerve. Voluntary muscle testing of the commonly examined peripheral nerves was done and graded as strong (S),weak (W), and paralyzed (P). Both W (weak) and P (paralyzed) was recorded as motor Nerve Function Impairment (NFI) present. For assessing hands: thumb up, little finger out and extension of the wrist against resistance was tested separately for both sides. Similarly for feet, dorsiflexion of feet against resistance was tested.

Any visible impairment on hands and feet like cracks/ wounds, absorption of fingers or toes, clawing of fingers or toes, contractures or any other impairment were looked for. For eyes, whether blinking of the eyes was Present (Pre) or absent (Abs) was noted Lid gap, if present, would be measured in mm by measuring scale and the patient’s ability to close the eyes, both lightly and tightly against resistance was also tested. Visual acuity was tested by a Snellen’s chart for each eye separately at 6 meters distance. WHO disability classification was followed for hands, feet and eyes. EHF (Eye, Hand and Feet) scores of an individual were calculated and would be ranged from 0 to 12. **^10^**

If skin lesion(s) with sensory loss was/were present, then the case was classified as indeterminate leprosy when there was only a hypopigmented macule, with no detection of nerve involvement (Paucibacillary or PB cases), or as one of the clinical forms defined by the Ridley-Jopling classification. **^11^** Slit skin smears for acid fast bacilli was done in all children with suspicious skin lesions. **^12^**

It was initially proposed that Lepromin test would be performed to detect subclinical infection in contacts, but was later decided against as many studies have suggested Lepromin test is inadequate to test subclinical infection with M. leprae.

## Results

Out of the 70 children brought under the study group, 41 were boys and 29 were girls *(Figure 1).* Boys formed 59% of the study group and girls 41%.

**Figure 1.**
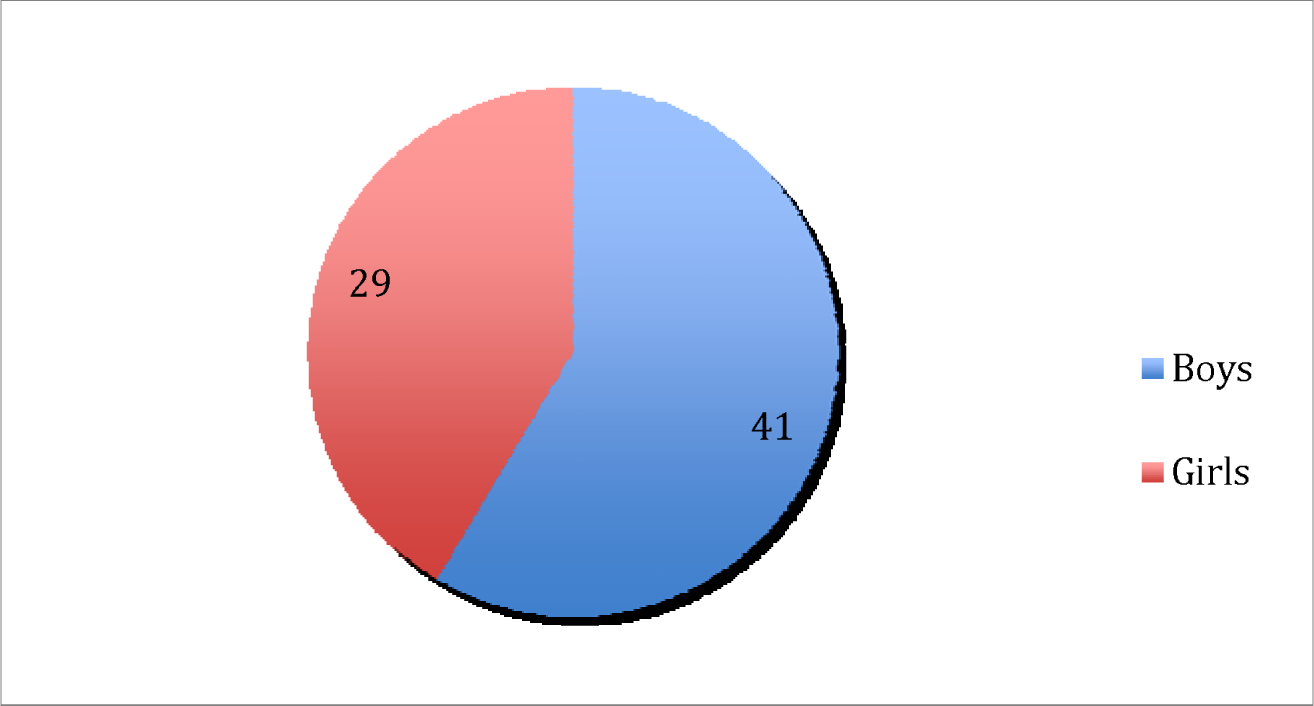
Sex Distribution.

The age distribution of the children is given in *Figure 2*. In 0-5 years, there were 10 children‐ 7 boys and 3 girls This group formed 14% of the study population. In 5-10 years age group were 16 children – 9 boys and 7 girls, forming 23% of the study group. Majority of the children belonged to the 10-15 years age group (44 children‐ 25 boys and 19 girls), contributing to 63% of the study population.

**Figure 2.**
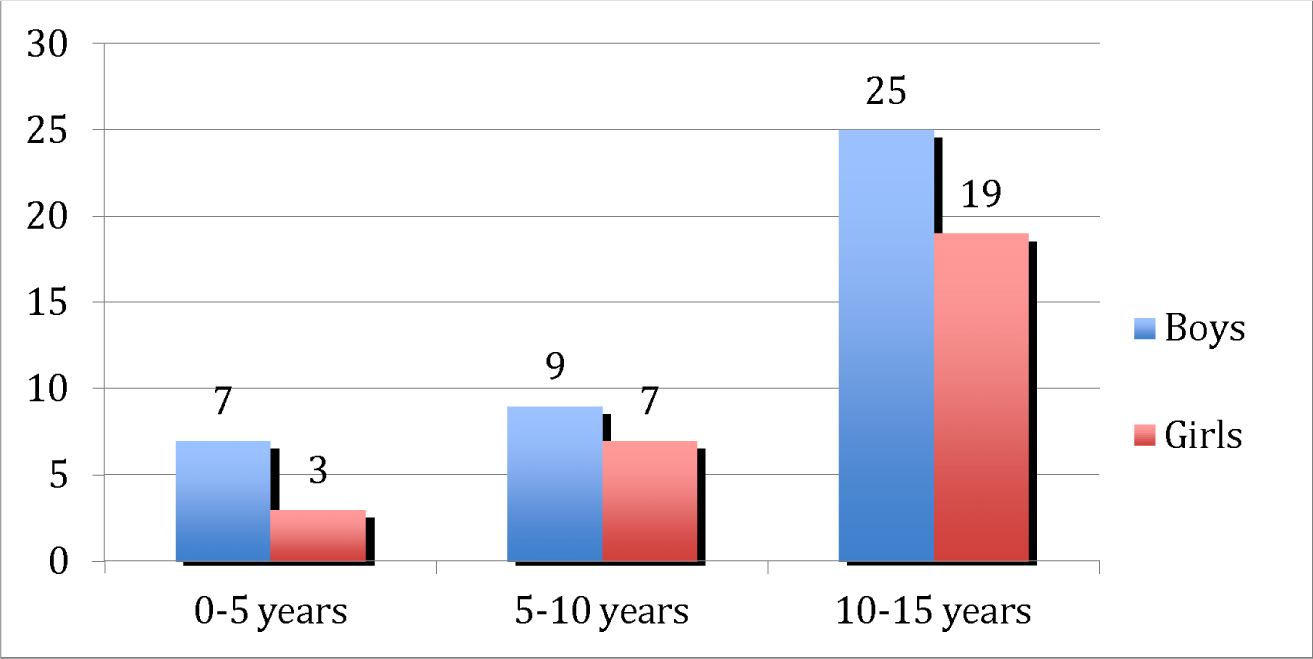
Age Distribution.

The anthropometric data obtained were compared with the respective WHO growth charts and the following results were obtained.

20 out of the 41 boys(49%) had heights which were in the Mean+/−2 SD range for their respective ages. 12 boys(29%) had heights which were between −2 and −3 SD for their respective ages, while 9 boys(22%) had heights which were below −3 SD for their respective ages. *(Figure 3)*

**Figure 3.**
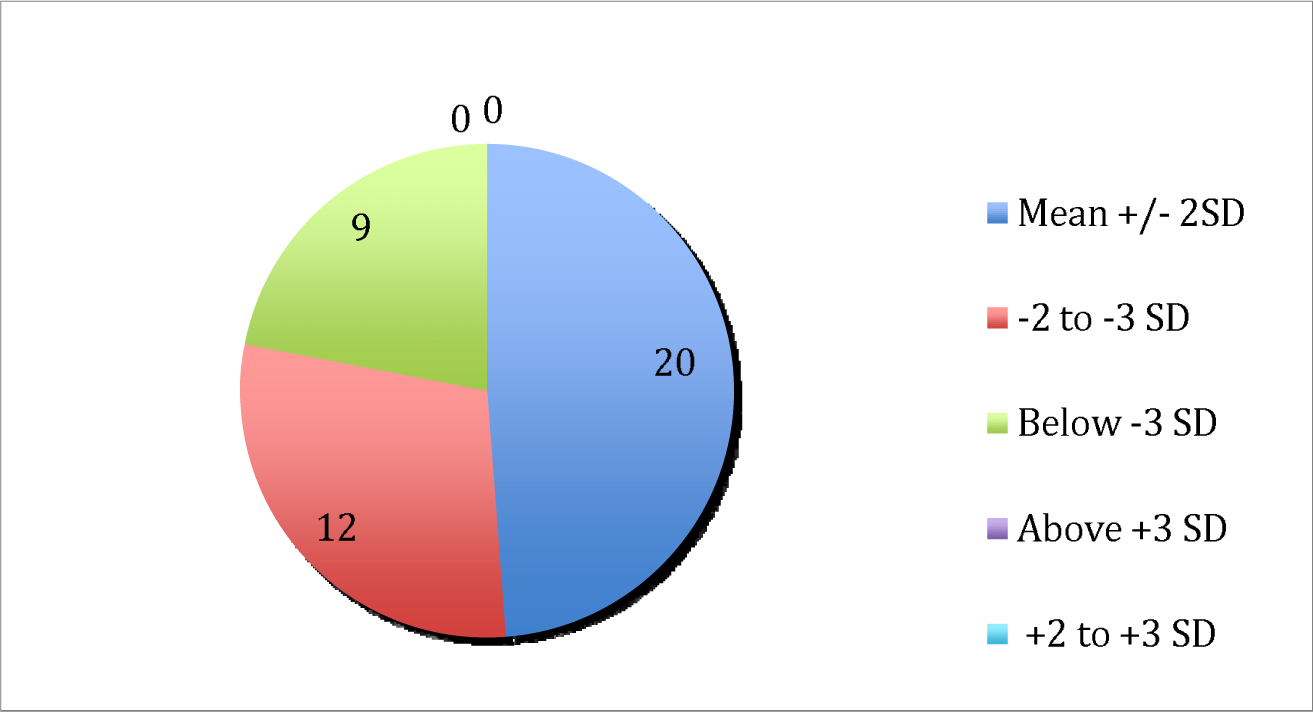
Height for age (boys)

20 out of 29 girls(69%) had heights which were in the Mean+/-2 SD range for their respective ages. 7 girls (24%) had heights which were between −2 and −3 SD for their respective ages, while 2 girls (7%) had heights which were below −3 SD for their respective ages. *(Figure 4)*

**Figure 4.**
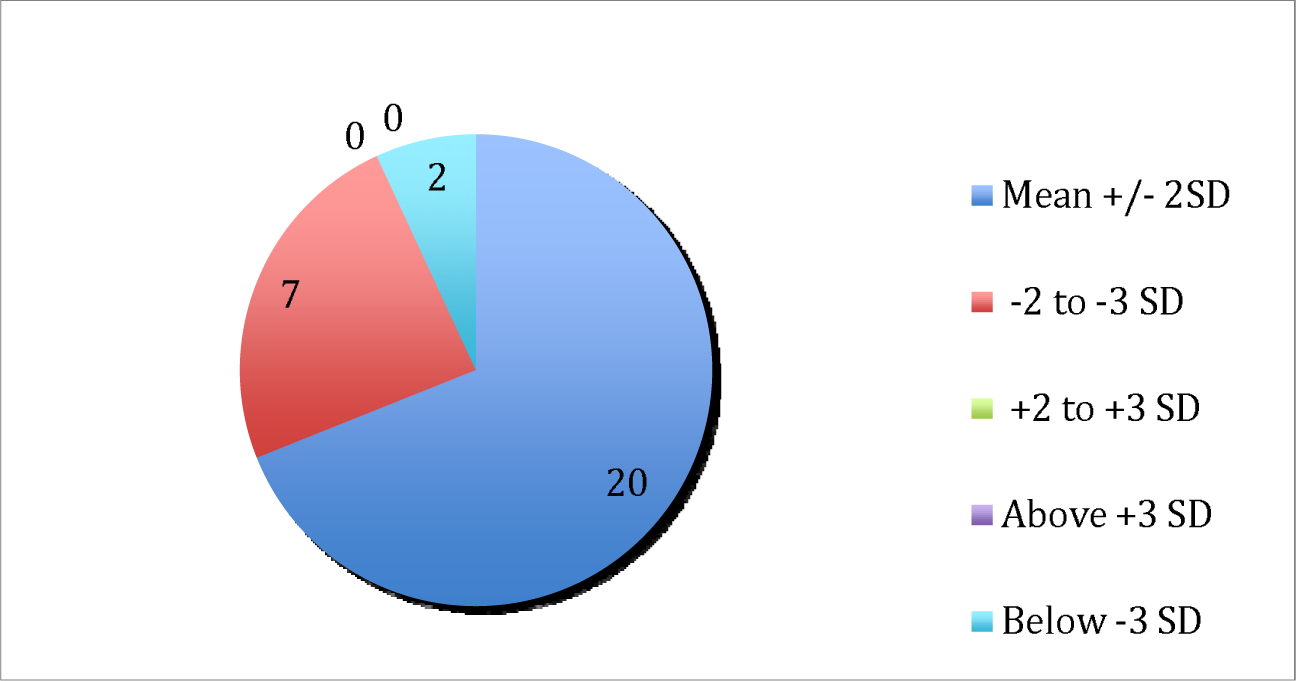
Height for age (Girls)

Out of the 10 children in the under 5 years age group(14% of the study group), 5 out of 7 boys had weight for height in Mean +/− 2SD range for their respective ages and 2 boys had weight for height between −2 and −3 SD for their respective ages, while all 3 girls under 5 years had weight for height in the Mean +/−2 SD for their respective ages.*(Figure 5)*

**Figure 5.**
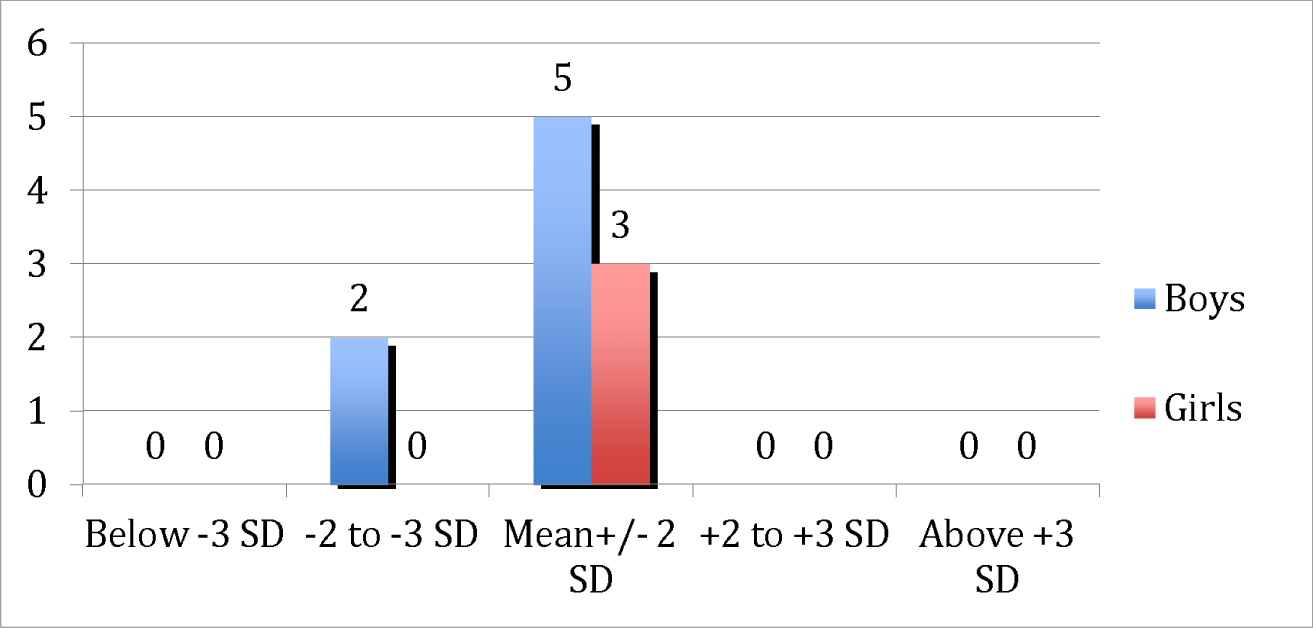
Weight for height (0-5 years)

26 children(37% of the study population) belonged to the 0-10 years age group – 16 boys and 10 girls. The weights of 8 out of 16 boys were in the Mean +/−2 SD range; 5 boys in between −2 and −3 SD and 3 boys below −3 Sd for their respective ages. Similarly among the 10 girls, weight of 7 girls were in the Mean +/−2 SD range and 3 girls below −3 SD for their respective ages.

Out of 41 boys, 19 had BMI in the Mean +/− 2 SD range, while 18 had BMI between −2 and −3 SD and 4 were below −3SD, with respect to the age-specific standards given in WHO growth charts. Among the 29 girls, 23 belonged to Mean +/−2 SD, 5 between −2 and −3 SD and 1 girl below −3 SD, with respect to age-specific standards. *(Figure 7)*

**Figure 6.**
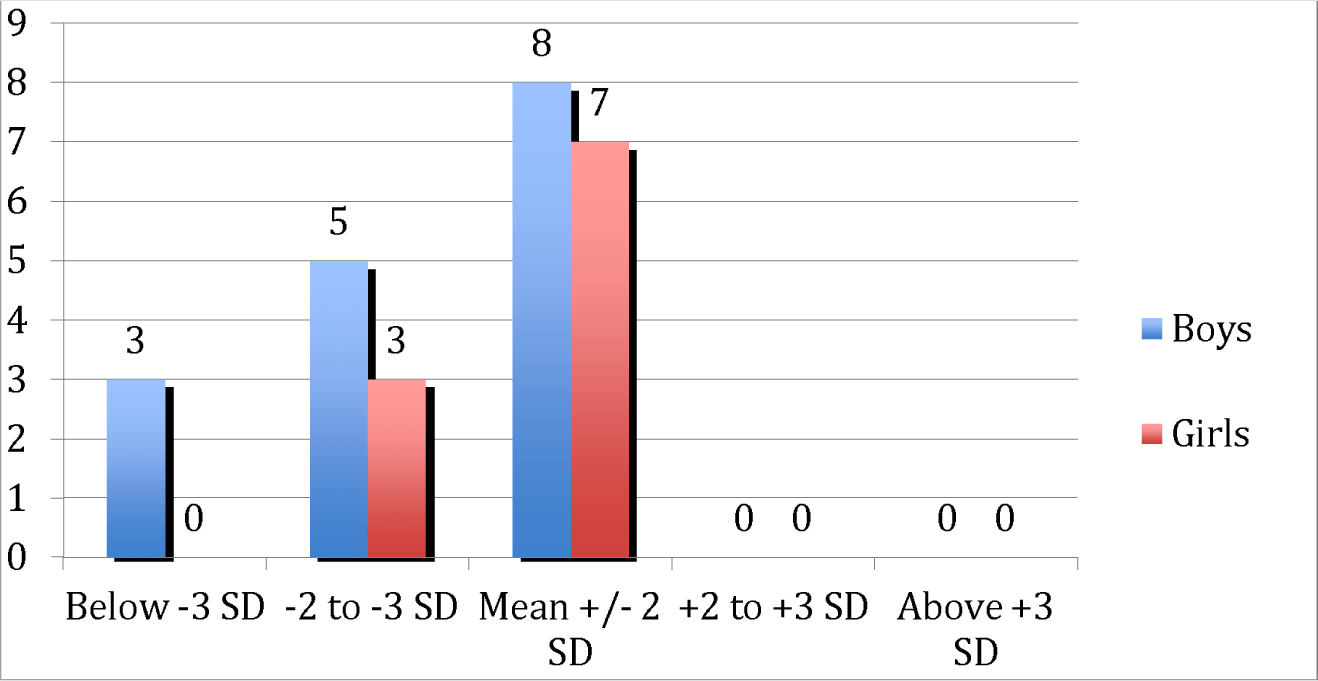
Weight for age (0-10 years)

**Figure 7.**
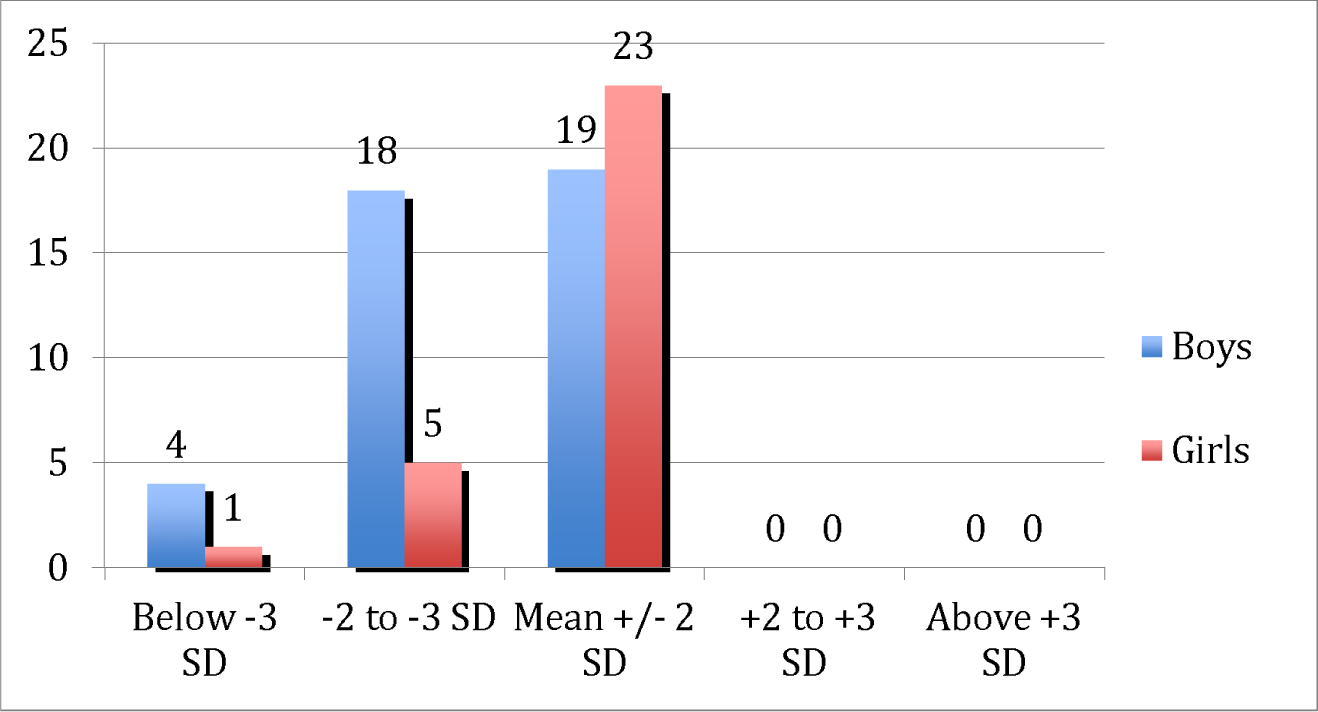
BMI for age.

Out of the 10 under 5 years children, 5 had head circumference in Mean +/-2 SD, 3 between −2 and −3 SD and 2 below −3 SD *(Figure 8).*

**Figure 8.**
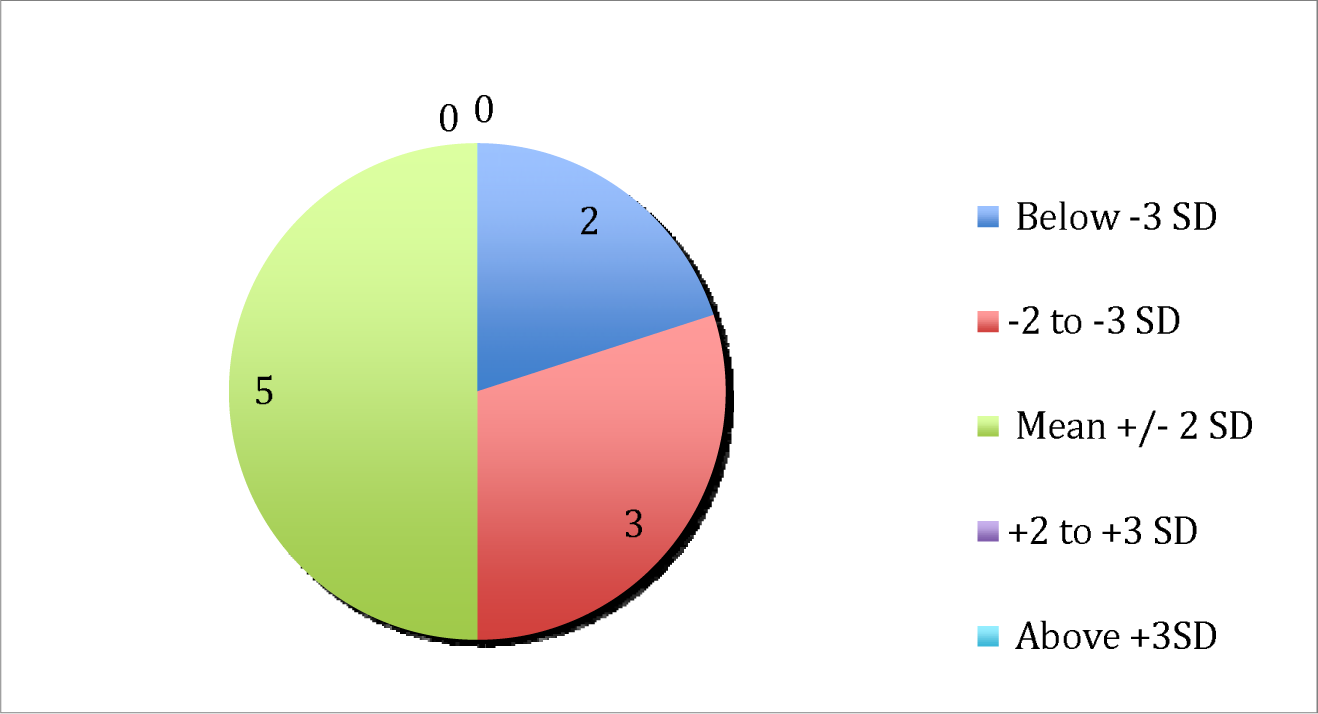
Head circumference (0-5 years)

All under 5 years children had Mid-arm circumference above 11.5 cm.

Socio economic status: According to Modified Kuppuswamy Scale, Out of 70 children assessed, 33 belonged to lower middle class (47%), 18 in the upper middle class (26%), 9 in the upper class (13%), 6 in the upper lower class (9%) and 4 in the lower class (5%) *(Figure 9).*

**Figure 9.**
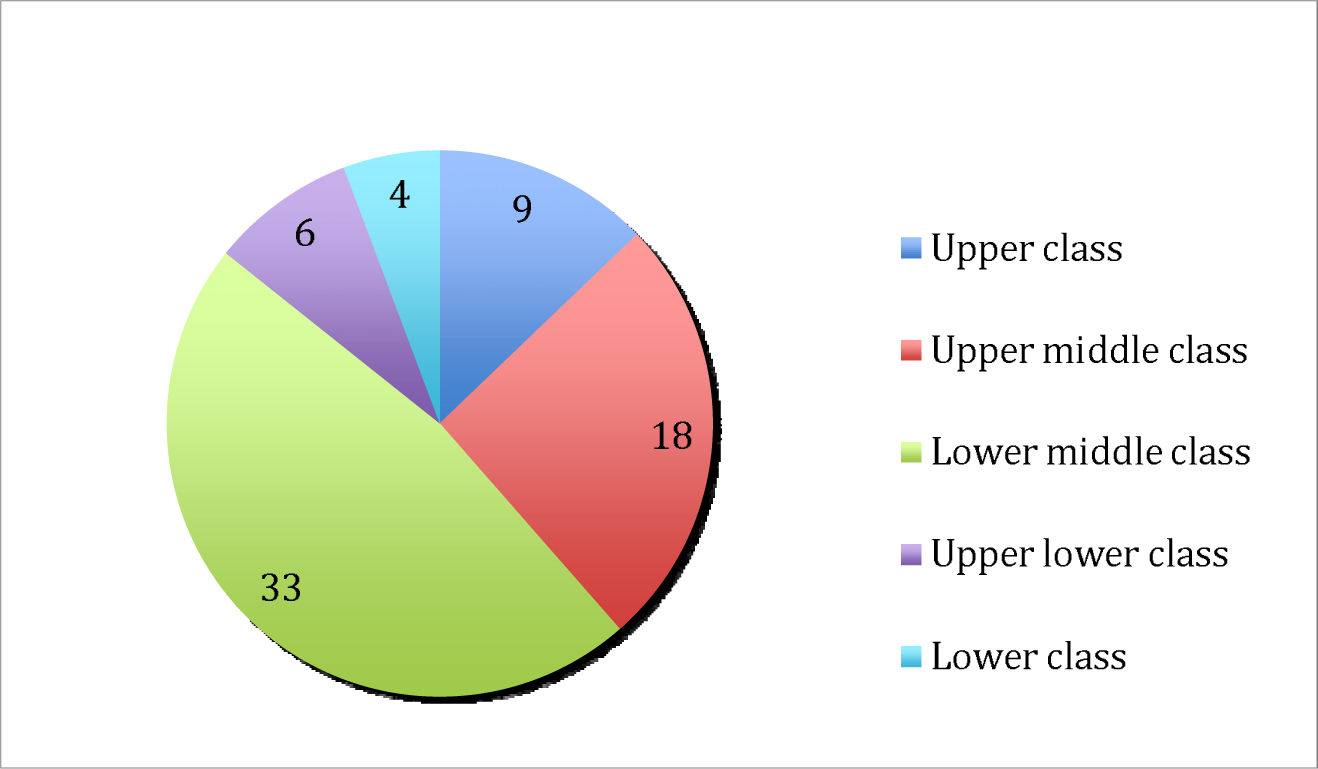
Socioeconomic status.

Household characteristics – From the responses to the questions related to household practices and characteristics, the following data were obtained. 40 children lived in overcrowded homes while 30 did not have overcrowding in their homes. Only 30 children lived in pucca houses while the remaining 40 children lived in semi pucca houses. All 70 children had access to latrines. 55 children had access to separate latrines for their houses, 15 children had to share latrines with other households. Only 9 children drank treated drinking water while 61 drank untreated drinking water. 64 out of 70 children washed hands regularly following different daily routine. 6 children did not wash hands that much regularly*(Figure 10). 64 children were completely immunized till date, while 6 of them were not.*

**Figure 10.**
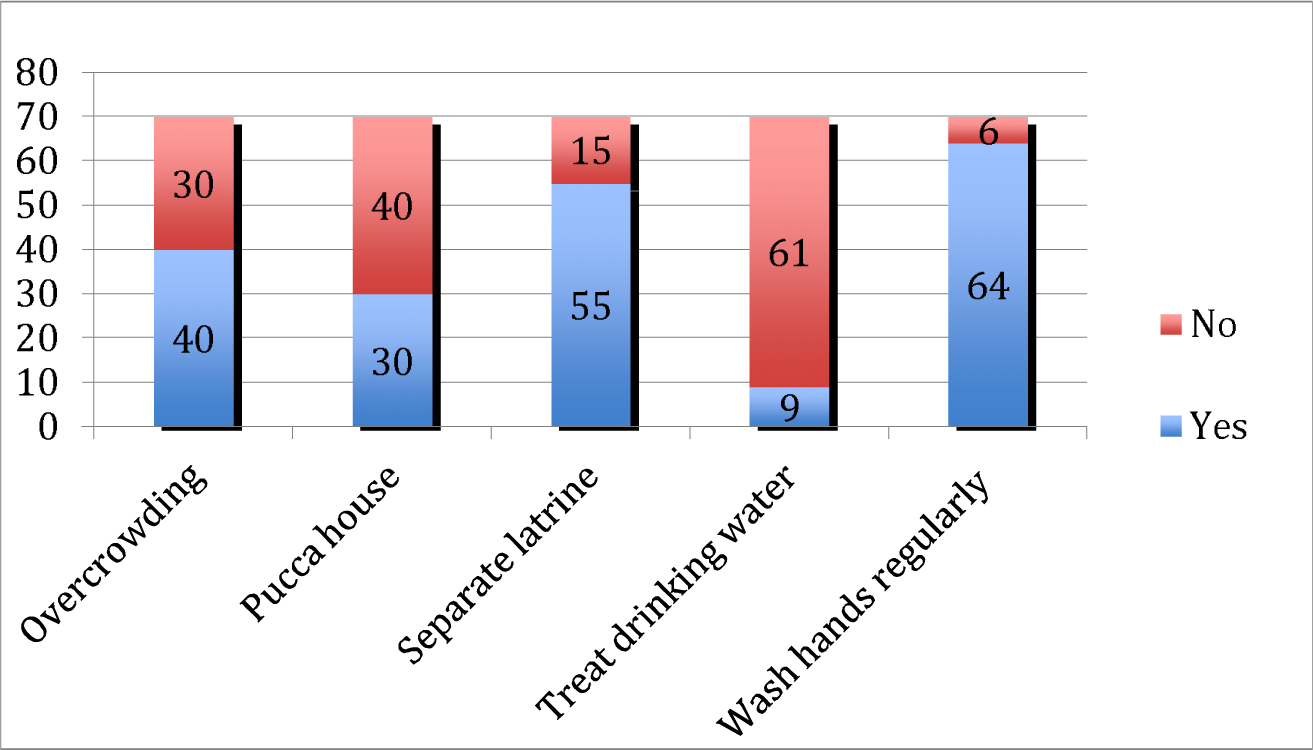
Household features.

Feeding practices – Based on the answers to the questions asked to the mother of the children, the following information was recorded. 61 children were breastfed immediately after birth, 6 within few hours after birth. 3 children were not breastfed at all. 30 children were given food other than breast milk for the first time before 6 months, 33 children at 6 months, 4 children between 6 and 12 months and 3 months at 1-2 years. 51 children received rice porridge as their first feed at weaning, while 19 children were fed small biscuits. Breastfeeding was stopped before 2 years for 58 children, at 2 years for 3 children, at 2-5 years for 6 children while 3 children were not breastfed at all*(Table 1)*.

**Table 1.** Feeding practices.

| Feeding practice |  |  | Observation(out of 70) |
| --- | --- | --- | --- |
| Start of breastfeeding | i. | Immediately after birth | 61 |
|  | ii. | Within few hours after birth | 6 |
|  | iii. | >24 hours after birth | 0 |
|  | iv. | Did not breastfeed at all | 3 |
| Start of Weaning | i. | <6 months | 30 |
|  | ii. | At 6 months | 33 |
|  | iii. | 6-12 months | 4 |
|  | iv. | 12- 23 months | 3 |
|  | v. | At or after 24 months | 0 |
| First food given | i. | Rice porridge | 51 |
|  | ii. | Rice powder | 0 |
|  | iii. | Small biscuit | 19 |
|  | iv. | Fruit juice | 0 |
| Stop of breastfeeding | i. | <2 years | 58 |
|  | ii. | At 2 years | 3 |
|  | iii. | 2-5 years | 6 |
|  | iv. | Not yet stopped | 0 |
|  | v. | Not breastfed | 3 |

**Table 2.** – Present or previous Co-morbidities.

| Present or previous co-morbidities | No. of children |
| --- | --- |
| Pallor | 19 |
| Past H/o exanthematous illnesses | 21 |
| Heat scald scars | 10 |
| Ventricular Septal Defect | 1 |
| Mental Retardation | 1 |
| Polymorphic Light Eruption | 5 |
| Umbilical hernia | 1 |

**Table 3.**
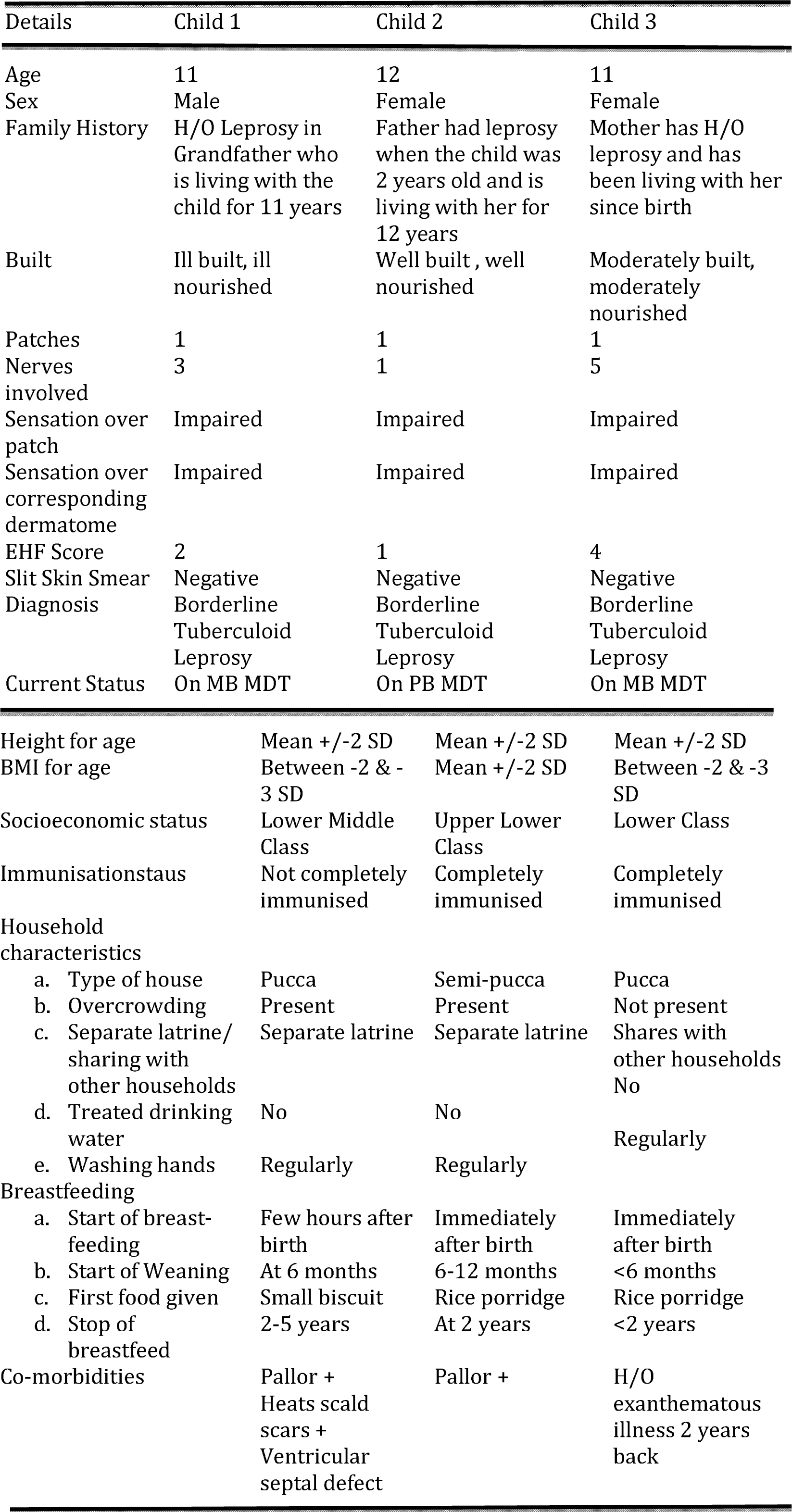
– Clinical Details of children diagnosed as leprosy.

The children’s mothers were asked about what food they most crave for and what food they avoid. 30 children craved for egg, while none avoided it. 12 children craved for fish and 3 avoided eating fish. 3 children avoided eating tubers while none craved for tubers. 3 children craved for Chicken while 9 children avoided it. 10 children avoided eating fruits while none craved for fruits. 25 children did not have any specific food craving while 45 children did not have any specific food they avoided *(Figure 11).*

**Figure 11.**
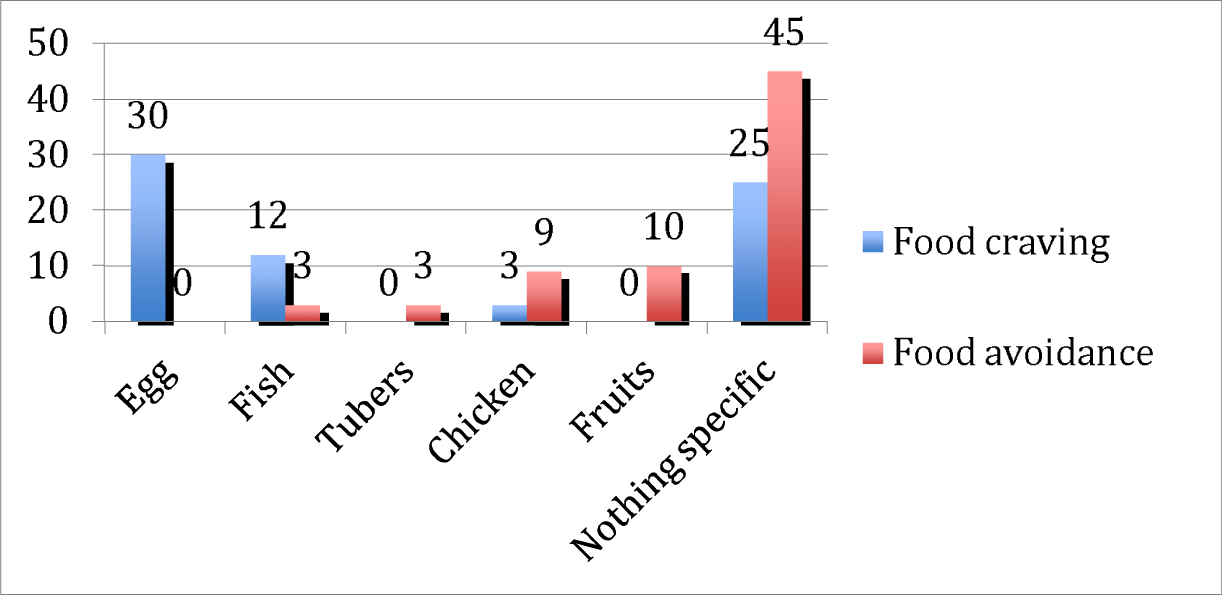
Food craving and food avoidance.

19 children had pallor. 21 children had recent or previous history of exanthematous illnesses. 10 children had heat scald scars present. 1 child was a known case of Ventricular Septal Defect and 1 child was a known case of Mental Retardation. 5 children were diagnosed to have polymorphic light eruption in clinical evaluation. 1 child had an umbilical swelling and then was further evaluated by the Department of Paediatrics at Chengalpattu Medical College Hospital and was diagnosed as a case of Umbilical hernia and is supposed to undergo surgery for the same.

5 children were found to have skin lesions suspicious of leprosy and were further evaluated at the Department of Dermatovenereoleprology at Chengalpattu Medical College Hospital. 3 children were diagnosed as leprosy while the other 2 children were found to have nutritional achromia and were given nutritional supplements for the same. The details of the 3 children are given below.

## Discussion

The analyses of all the data collected have given greater insight into children with contact to leprosy in the rural community. In terms of sex distribution, boys:girls ratio is approximately 60:40. And a majority of the children fall under the 10-15 years age group, with all 3 children diagnosed as a case of leprosy through this study falling under the same age group.

After analyzing all the anthropometric factors, it is understood that girls have better anthropometric measurements compared to the boys. This can be substantiated as follows:

According to WHO, the normal range of values for anthropometric measurements is the Mean+/−2 SD(−2 to +2 Z scores; SD scores are also known as Z scores) range and 95% of a distribution fall in this range. Height for age, Weight for age and Weight for height Z scores falling between −2 and −3 SD indicate moderate malnutrition, and scores below −3 SD indicate severe malnutrition and a score of more than +3 SD indicates obesity. **^13^**

9 boys and 2 girls have height for age below −3 SD, 4 boys and 1 girl have BMI for age below −3 SD and 3 boys with weight for age below −3 SD all indicating severe malnutrition. This shows 7-22% of boys are severely malnourished, while only 3-6% of girls fall under this category. Overall, out of all 70 children, 4-15% children are severely malnourished.

18 boys and 5 girls have BMI for age between −2 and −3 SD, 12 boys and 7 girls have height for age between −2 and −3 SD, 5 boys and 3 girls have weight for age between −2 and −3 SD and 2 boys have weight for height between −2 and −3 SD, implying that 5-44% of the boys are moderately malnourished, while only 10-24% of the girls belong to this category. Overall, 3-33% of the children are moderately malnourished. Combining both, 3-48% of the children are malnourished, irrespective of the type of malnourished. None of the children assessed are overweight or obese.

40 of the 70 children live in overcrowded semi-pucca houses. 61 children drink untreated drinking water. This can be probably attributed to 30% of the children having history of exanthematous illnesses. 64 children wash their hands regularly implying most of the children have good personal hygiene.

Most(61) of these children were breastfed immediately after birth, 6 within few hours after birth and 3 children were not breastfed at all. 30 children were weaned before 6 months, 33 at 6 months, 4 after 6 months and 3 children were started on foods only after 1 year, which is significant. Most children (51) were given rice porridge as first feed and some (19) were given small biscuits. Breastfeeding was stopped before 2 years in 58 children, at 2 years in 3 children and between 2 and 5 years in 6 children and as stated above 3 children were not breastfed at all.

It is to be noted that breastfeeding is an important factor to prevent malnutrition and infection in children. Introduction of complementary feeds at 6 months is also equally important because they provide for the increased nutrient needs of the children after 6 months. Moreover, if feeds other than breast milk are not started at 6 months, then it might be difficult for children to accept foods with new tastes and textures in the future. From the data collected, it can be implied that about 30-35 children fall under the malnourished category. 30 children were exclusively breastfed for less than 6 months. So this could also have attributed to the malnutrition in these children. **^14^**

19 out of the 70 children had clinical pallor. Presence of pallor is a reliable indicator of anemia. The World Health Organisation (WHO) estimates that 25% of pre schoolers and 40% of school children are anemic. All the 19 children belonged to the school going age group. They formed 27% of the study population. This is lesser compared to the WHO estimate.**^15^**

As per the National Leprosy Eradication Programme Annual Report 2015-2016, Children formed 8.94% of the new cases detected throughout the country during 2015-16. The child case rate in Tamil Nadu is 15.86%, out of which 1.50% were Multibacillary cases and 14.36% Paucibacillary cases. No. of new child with Grade 2 disability for Tamil Nadu was 1. **^16^**

Among the 70 leprosy contact children, 3 children were diagnosed to have leprosy. This implies 4.28% of the children assessed had leprosy.This can be attributed the high prevalence of leprosy in Chengalpattu and the fact that all childen were leprosy contacts. Of the 3, 2 children had multibacillary leprosy while 1 had paucibacillary leprosy, according to the WHO classification. **^17^** All 3 children were classified as cases of Borederline Tuberculoid Leprosy according to the Ridley-Jopling classification.**^18^** There isn’t accurate statistical data available as to how many under 15 years old children are present in Chengalpattu. Hence the new child case detection rate for Chengalpattu could not be calculated.

All these 3 children had contact to leprosy for 10 or more years living with them, before being diagnosed. According to WHO, the average incubation period for M. leprae is 5 years. It can range from about 1 year to as long as about 20 years. So all these 3 children fall under this range, fitting with the long incubation period of leprosy. **^19^** This study reflected the same observation, that closeness of contact and duration of contact to leprosy is a significant risk factor, which was also reflectedin various childhood contact studies that were reviewed.

Child 1, who was incompletely immunized, was a previously diagnosed case of congenital heart disease (ventricular septal defect). There is no evidence generated yet to correlate congenital heart disease and leprosy; further research is needed in this research. 2 out of the 3 children diagnosed as leprosy were malnourished (66.6%). So it is evident that malnutrition can be a risk factor for leprosy in children who are close contacts.

There has also been a significant amount of exanthematous illness in this group of children. It’s not known whether exanthematous illness can be a precipitating factor for leprosy. This is also an area which needs further research.

## Conclusion

Through this study it can be concluded that,

- Malnutrition is a significant risk factor for leprosy.
- The closeness and duration of contact to leprosy is an important risk factor.
- Incomplete immunization in combination with the other risk factors is morbidity‐ inducing.
- Regular contact screening and early case detection are essential strategies to prevent further transmission in the endemic areas.
- Diagnostic methods for detection of subclinical infection in contacts needs further research.

This study was conducted only for a small group of 70 children living in a highly endemic area (Chengalpattu) with contact to leprosy. The results obtained from this study donot reflect the whole population. Hence further large scale studies regarding the nutritional status and morbidity of such children must be carried out to gain results reflecting the whole population. Future research in leprosy needs to be focused on child contacts to leprosy. There have to be measures taken to implement contact tracing on a large scale as part of the leprosy control programmes.

